## Supporting Information for "Optimal evaluation of energy yield and driving force in microbial metabolic pathway variants"

#### S1. Acid-Base Equilibria

The pH-based speciation of acidic species occurs at a timescale that is several orders of magnitude faster than metabolic reactions, and thus it is typical to assume those species to be at equilibrium. This implies that the protonated form of a species and all its successive deprotonated forms are at the same chemical potential, and thus the choice of microspecies does not affect the rest of the framework constraints. The underlying assumption in such a situation is that the bounds described in section 2.5 are applied to the total concentration of each species rather than individual microspecies.

However, it may be desired in some models to select one species as the true substrate of the metabolic reactions and apply the concentration bounds to that form. Furthermore, different reactions in the same pathway may use two different forms as the true substrate, meaning the two species concentrations have to be included as decision variables, but are not independent of one another. An example of this is the dicarboxylate-4-hydroxybutyrate cycle in autotrophic $CO_2$ fixing archaea, where both $CO_2$ and $HCO_3^-$ are true substrates for different reactions in the pathway [1]. In such a case, it becomes necessary to introduce an equality constrain based on the definition of acid dissociation constant:

$$K_a = \frac{C_{H^+} * C_{A^-}}{C_{HA}} \tag{1}$$

where HA is the protonated form of a species, $A^-$ is its deprotonated form, and $K_a$ is the acid dissociation constant associated with the deprotonation. Fortunately, by applying the logarithm to equation (17), we can obtain a linear equality constrain:

$$\ln C_{A^-} - \ln C_{HA} = \ln 10 * (pH - pK_a) \tag{2}$$

### S2. Case study pathway information

The following reaction stoichiometries, possibility of proton translocation and permissible electron carriers have been assembled using previous work [2] and biochemical databases [3,4]. Note that the molar ratio of the electron carrier regeneration reactions vary depending on the variant being evaluated.

**Table S1**: Reaction and biochemical information for propionate oxidation pathways. The numbering of the pathways is as presented in the figure in the main text.

| Reaction | Molar Ratio | | | | | | SLP | Prot trans allowed? | Electron Carriers |
|---|---|---|---|---|---|---|---|---|---|
| | P1 | P2 | P3 | P4 | P5 | P6 | | | |
| Propanoic Transport (In) | 1 | 1 | 1 | 1 | 1 | 1 | 0 | Yes | |
| Propionate -> Propionyl-CoA | 1 | 1 | 0 | 0 | 1 | 0 | -1 | No | |
| Propionate + Acetyl-CoA -> Acetate + Propionyl-CoA | 0 | 0 | 1 | 0 | 0 | 1 | 0 | No | |
| Propionyl-CoA -> methylMalonyl-CoA | 1 | 0 | 1 | 0 | 0 | 0 | -1 | Yes | |
| Propionyl-CoA + Oxaloacetate -> methylMalonyl-CoA + Pyruvate | 0 | 1 | 0 | 0 | 0 | 0 | 0 | No | |
| mMalon-CoA -> Succinyl-CoA | 1 | 1 | 1 | 0 | 0 | 0 | 0 | No | |
| Succinyl-CoA -> Succinate | 1 | 1 | 1 | 0 | 0 | 0 | 1 | No | |
| Succinate -> Fumarate | 1 | 1 | 1 | 0 | 0 | 0 | 0 | Yes | FAD, Quinone |
| Fumarate -> Malate | 1 | 1 | 1 | 0 | 0 | 0 | 0 | No | |
| Malate -> Oxaloacetate | 1 | 1 | 1 | 0 | 0 | 0 | 0 | No | NAD, FAD |
| Oxaloacetate -> Pyruvate | 1 | 0 | 1 | 0 | 0 | 0 | 1 | Yes | |
| Pyruvate -> Acetyl-CoA | 1 | 1 | 1 | 1 | 0 | 0 | 0 | No | Fd |
| Propionate + Lactoyl-CoA -> Lactate + Propionyl-CoA | 0 | 0 | 0 | 1 | 0 | 0 | 0 | No | |
| Propionyl-CoA -> Acryloyl-CoA | 0 | 0 | 0 | 1 | 1 | 1 | 0 | Yes | FAD, Quinone |
| Acryloyl-CoA -> Lactoyl-CoA | 0 | 0 | 0 | 1 | 0 | 0 | 0 | No | |
| Lactate -> Pyruvate | 0 | 0 | 0 | 1 | 0 | 0 | 0 | No | NAD |
| Acryloyl-CoA -> 3-hydroxyPropionyl-CoA | 0 | 0 | 0 | 0 | 1 | 1 | 0 | No | |
| 3-hydroxyPropionyl-CoA -> 3-hydroxyPropionate | 0 | 0 | 0 | 0 | 1 | 1 | 1 | No | |
| 3-hydroxypropionate -> Malonate semialdehyde | 0 | 0 | 0 | 0 | 1 | 1 | 0 | No | NAD |
| Malonate semialdehyde -> Acetyl-CoA | 0 | 0 | 0 | 0 | 1 | 1 | 0 | No | NADP |
| Acetyl-CoA -> Acetate | 1 | 1 | 0 | 1 | 1 | 0 | 1 | No | |
| Acetate Transport (Out) | 1 | 1 | 1 | 1 | 1 | 1 | 0 | Yes | |
| $FADH_2 + NAD^+ \rightarrow FAD^{2+} + NADH$ | | | | | | | 0 | Yes | |
| $UQ_{red} + NAD^+ \rightarrow UQ_{ox} + NADH$ | | | | | | | 0 | Yes | |
| $Fd_{red} \rightarrow Fd_{ox} + H_2$ | | | | | | | 0 | Yes | |
| $NADH \rightarrow NAD^+ + H_2$ | | | | | | | 0 | Yes | |
| $NADPH \rightarrow NADP^+ + H_2$ | | | | | | | 0 | Yes | |

**Table S2**: Reaction and biochemical information for the reverse TCA cycle pathway. Note that the molar ratios for the electron carrier reactions will be negative, indicating that they will proceed in the opposite direction to the one listed in the table

| Reaction | Molar Ratio | SLP | Prot trans allowed? | Electron Carriers |
|---|---|---|---|---|
| $CO_2$ Transport | 2 | 0 | Yes | |
| Succinyl CoA -> 2-ketoglutarate | 1 | 0 | No | NAD, NADP, Fd |
| 2-ketoglutarate -> Isocitrate | 1 | 0 | Yes | NAD, NADP |
| Isocitrate -> Aconitate | 1 | 0 | No | |
| Aconitate -> Citrate | 1 | 0 | No | |
| Citrate -> Oxaloacetate + Acetyl CoA | 1 | -1 | No | |
| Oxaloacetate -> Malate | 1 | 0 | Yes | NAD, NADP, Quinone |
| Malate -> Fumarate | 1 | 0 | No | |
| Fumarate -> Succinate | 1 | 0 | Yes | NAD, FAD, Quinone |
| Succinate -> Succinyl CoA | 1 | -1 | No | |
| $FADH_2 + NAD^+$ -> $FAD^{2+}$ + NADH | | 0 | Yes | |
| $UQ_{red} + NAD^+$ -> $UQ_{ox}$ + NADH | | 0 | Yes | |
| $Fd_{red}$ -> $Fd_{ox} + H_2$ | | 0 | Yes | |
| NADH -> $NAD^+ + H_2$ | | 0 | Yes | |
| NADPH -> $NADP^+ + H_2$ | | 0 | Yes | |

### S4. Thermodynamic information

**Table S3**: Standard enthalpies and Gibbs energy changes of formation for species used in this work

| Species Name | $\Delta G^0_f$ (kJ/mol) | $\Delta H^0_f$ (kJ/mol) | Source |
|---|---|---|---|
| Propionate | -361.08 | -510.4 | [2] |
| Succinyl CoA | -461.89 | -619.32 | [5] |
| Coenzyme A | 0 | 0 | [5] |
| Acetyl CoA | -140.67 | -227.77 | |
| 2-ketoglutarate | -793.41 | -1044.1 | [5] |
| iso-citrate | -1156 | N/A | [5] |
| cis-Aconitate | -917.13 | N/A | [5] |
| Citrate | -1162.7 | -1515.1 | [5] |
| Oxaloacetate | -793.82 | -959.9 | [5] |
| Malate | -843.22 | -1079.8 | [6] |
| Fumarate | -602.27 | -777.91 | [6] |
| Succinate | -690.91 | -909.29 | [7] |
| Pyruvate | -472.27 | -596.22 | [6] |
| Malonate semialdehyde | -477.6 | -609.39 | [5] |
| 3-hydroxypropionate | -518.4 | -676.04 | [5] |
| Hydroxypropionyl-CoA | -285 | -400.64 | [5] |

| | | | |
|---|---|---|---|
| Acryloyl CoA | -48.5 | -122.98 | [8] |
| Propionyl CoA | -131.31 | -248.41 | [5] |
| Methylmalonyl CoA | -454.65 | -624.6 | [5] |
| Lactate | -517.81 | -687 | [2] |
| Lactoyl CoA | -287.8 | -405.92 | [8] |
| Hydrogen | 17.55 | -4.16 | [8] |
| Acetate | -369.41 | -486 | [2] |
| Ubiquinone (ox) | 0 | N/A | [5] |
| Ubiquinone (red) | -64.43 | N/A | [5] |
| Ferredoxin (ox) | 0 | N/A | [7] |
| Ferredoxin (red) | 79.1177 | N/A | [7] |
| FAD | 0 | N/A | [7] |
| FADH2 | -37.4 | N/A | [7] |
| NAD+ | 0 | N/A | [2] |
| NADH | 21.83 | N/A | [2] |
| NADP+ | 0 | NA | [2] |
| NADPH | 21.83 | NA | [2] |
| Carbon Dioxide | -386 | -413.8 | [2] |
| Proton | 0 | 0 | |
| Water | -237.18 | -285.8 | [5] |

### S5. Multiplicity of solutions to the optimization problem

As discussed in section 3.1.1 of the main text, the concentration profiles and configuration of proton translocations obtained by the solvers for an optimized pathway variant is not unique. Thus, there are several profiles that lead to the same optimum energy recovery and driving force distribution. In the figures below, we present multiple such profiles for the cyclical hydroxypropionyl pathway.

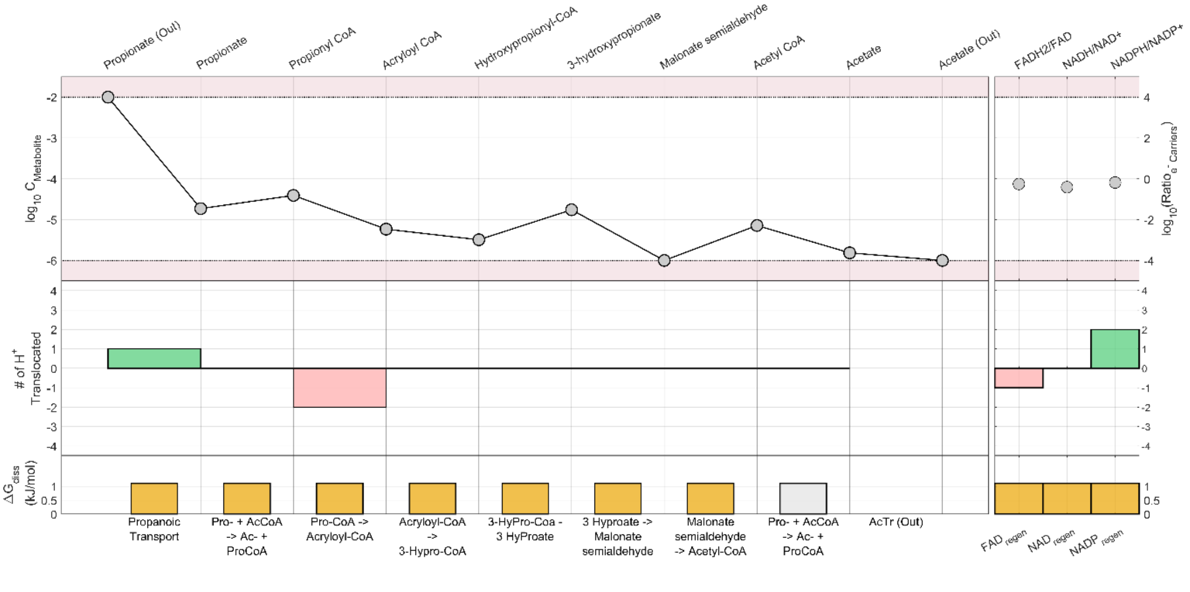

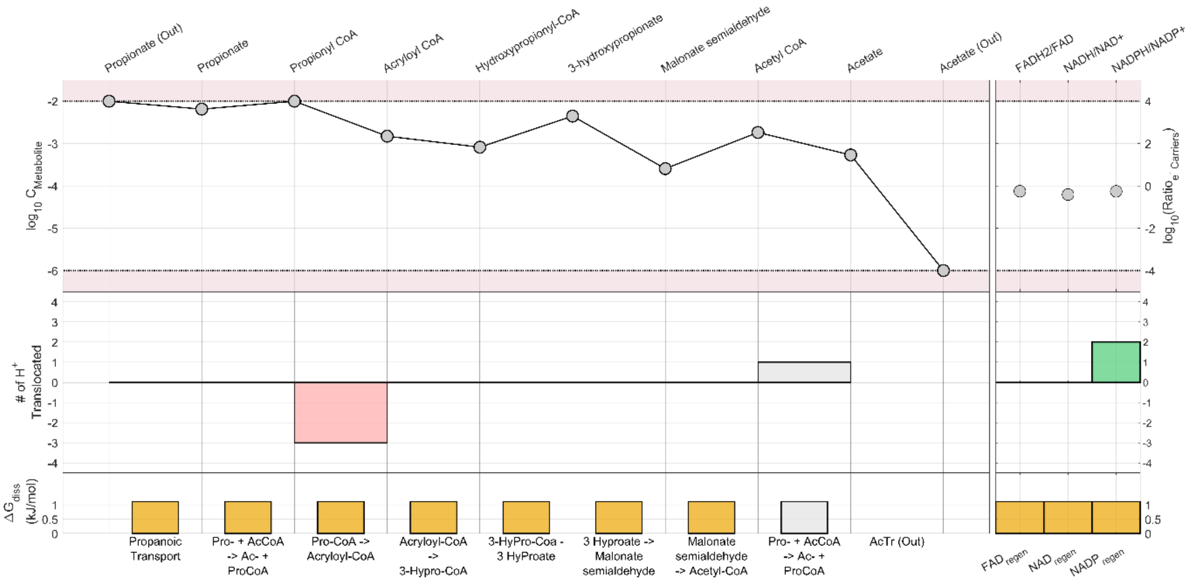
